# Kynurenic acid promotes activity-dependent synaptic pruning in schizophrenia

**DOI:** 10.1101/2023.10.19.563090

**Authors:** Funda Orhan, Susmita Malwade, Neda Khanlarkhani, Asimenia Gkoga, Oscar Jungholm, Marja Koskuvi, Šárka Lehtonen, Lilly Schwieler, Kent Jardemark, Jari Tiihonen, Jari Koistinaho, Sophie Erhardt, Göran Engberg, Samudyata Samudyata, Carl M. Sellgren

## Abstract

Schizophrenia is a neurodevelopmental disorder characterized by an excessive loss of synapses. Recent data suggest that this is due to increased microglia-mediated synaptic pruning. Here, we utilize human induced pluripotent stem cell-derived models to show that kynurenic acid (KYNA), an endogenous NMDA-receptor antagonist observed to be increased in the brains of individuals with schizophrenia, reduces neuronal activity and promote microglial uptake of synapses. In a human brain organoid model, we confirm reduced microglia-mediated synaptic pruning upon inhibiting the endogenous KYNA production. To verify our experimental data in a clinical context, we integrate large-scale transcriptomic and genetic datasets and show that KYNA-producing kynurenine aminotransferases (KATs) enrich for genes governing synaptic activity and genetic risk variants for schizophrenia. Together, these results link genetic risk variants for schizophrenia to elevated production of KYNA and excessive activity-dependent synaptic pruning, while implicating pharmacological inhibition of KATs as a strategy to avoid synapse loss in schizophrenia.

## Introduction

Schizophrenia (SCZ) is a highly heritable and devastating psychiatric disorder with a genetic architecture characterized by a prominent polygenic component.^1^ Large-scale genome-wide association studies (GWAS) have successfully uncovered a growing number of common genetic variants linked to increased SCZ risk,^2^ but any biological insight regarding these risk variants is still sparse. One exception is the complement component 4 (*C4*) locus, in which repeated copy numbers of *C4A* increase SCZ risk as well as the extent of microglia-mediated synaptic pruning.^3,4^ These findings align with a reduced spine density in SCZ^5–7^, but only partly explain the excessive synaptic elimination in SCZ models.^4^ One plausible explanation is the contribution of synaptic pathology, and animal studies have indicated that complement dependent microglial engulfment of synapses is activity-dependent such that synapses with decreased activity are more likely to be tagged by complement factors.^3,8,9^ In line with this, genes prioritised within GWAS associated loci by fine-mapping, then also enriching for rare risk variants, largely converge on pathways governing synapse biology. ^2^

This prompted us to consider a role of kynurenic acid (KYNA) in amplifying pathological synaptic pruning in SCZ. KYNA is a neuroactive metabolite of tryptophan along the kynurenine pathway and inhibit all three ionotropic glutamate receptors at millimolar concentrations.^10^ However, in the mammalian brain, local KYNA concentrations at the synapse can be expected to be in the nanomolar to low micromolar range, suggesting that the targets of endogenous KYNA are restricted to the obligatory glycine co-agonist site of the *N*-methyl-D-aspartate receptor (NMDAR), and possibly the α7 nicotinic acetylcholine receptor (α7nAchR) receptor.^10^ Within this concentration range, an elevation of KYNA levels has also been shown to induce SCZ-related phenotypes in rodents such as disruption in pre-pulse inhibition,^11^ increased striatal dopamine release by amphetamine,^12^ and an augmentation of the amphetamine-induced locomotor response.^13^ Further, clinical studies have also repeatedly reported elevated levels of KYNA in CSF ^14–17^ as well as in postmortem brain of individuals with SCZ.^18,19^

To address this question, we first generated excitatory neurons from human induced pluripotent stem cells (iPSCs) and confirmed that KYNA decreases the activity of excitatory neurons. After adding human microglia-like cells, KYNA also caused a decrease in spine density which was reversed by adding NMDA and the co-agonist D-serine. Utilizing a forebrain organoid model showing spontaneous microglial synaptic pruning, we then inhibited the innate production of KYNA and observed a decrease in microglial uptake of synaptic structures. To address the clinical relevance of these experimental findings, we integrated large-scale postmortem brain tissue and genetic datasets to show that the gene co-expression networks for the KYNA-producing enzymes, i.e., the kynurenine aminotransferases (KATs), are independently enriched for genes governing synaptic activity (including the genes coding for the NMDAR subunits and the α7nAchR) as well as common genetic risk variants for SCZ.

## Results

### KYNA decreases activity in human excitatory neurons

To confirm a human model in which KYNA could be used to manipulate the activity of excitatory neurons, we first used TALEN-based plasmids to insert a doxycycline-inducible *neurogenin 2* (NGN2) construct into the adeno-associated virus integration site 1 (*AAVS1*) safe-harbor locus of human iPSC-derived neural progenitor cells (NPCs) (Figure 1A).^4^ After transfection, NPCs were expanded to generate human cortical excitatory neuronal cultures (Figures 1B-D) and treated for 24 hours with either 25 or 50 μM KYNA after which spontaneous calcium transients were monitored. We then observed a dose-dependent decrease in calcium transients, confirming the previously observed effect in the millimolar range in human iPSC-derived neurons^20^ (Figure 1E).

**Figure 1.**
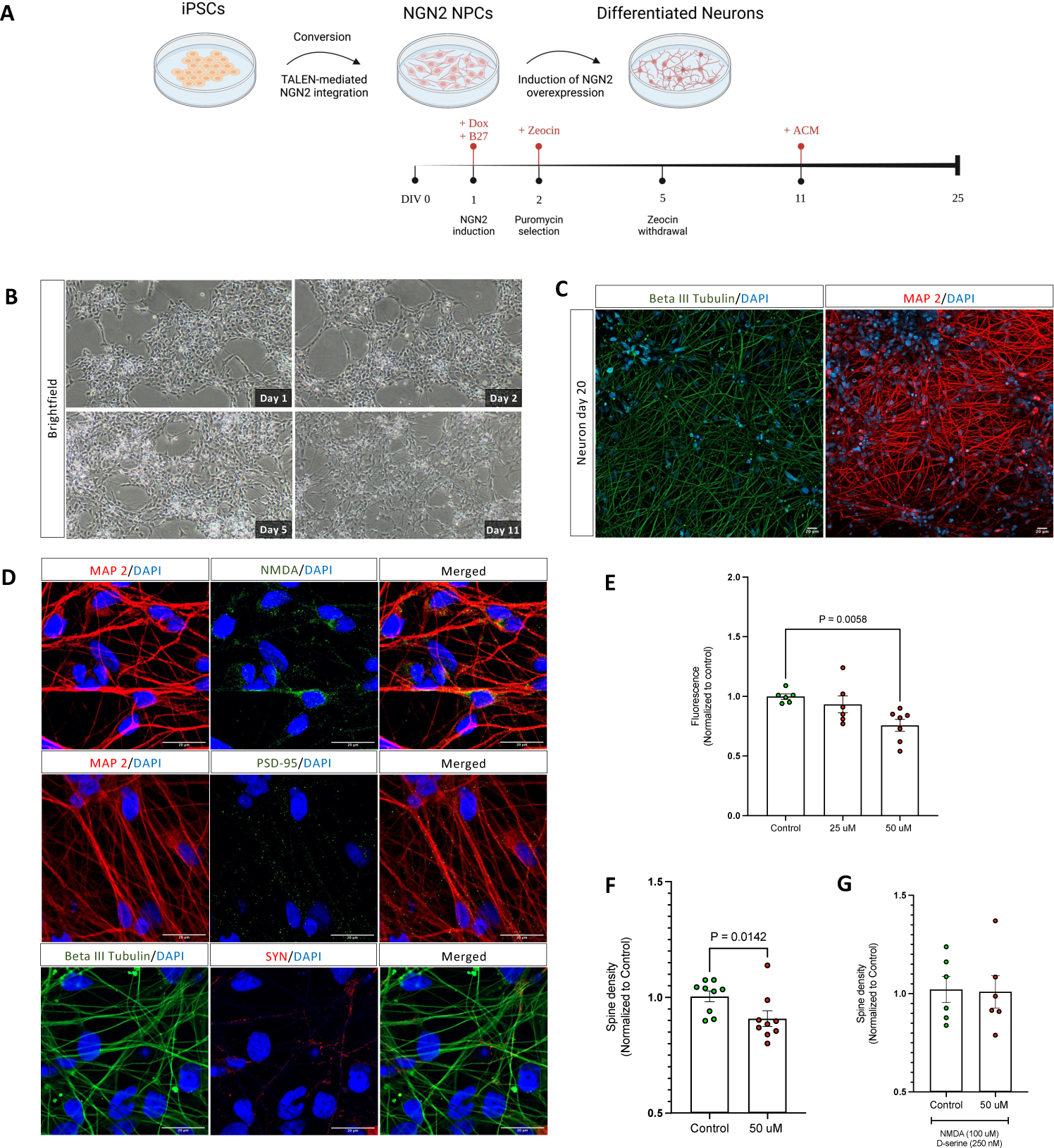
Kynurenic acid (KYNA) induces an activity-dependent decrease in spine density in human neuron-microglia co-cultures. (A) Schematic of the methodology used to generate iPSC-derived excitatory neurons. (B) Morphology of *NGN2*-induced excitatory neurons at various time points of induction. By day 20 of culture, the expression of (C) neuronal and (D) synaptic markers was confirmed by immunofluorescence with specific antibodies against Beta III Tubulin, Microtubule-Associated Protein 2 (MAP2), NMDA, postsynaptic density protein 95 (PSD-95), and Synapsin 1 (SYN). Confocal images were captured at 20x (C) and 63x (Airyscan) (D). Scale bars for the representative images are 20 μm. (E) 24 hours pre-treatment with KYNA induced a dose-dependent effect on intracellular calcium levels in cultures of human excitatory neurons. The experiment was performed on day 23 neurons in triplicate and repeated twice. F) Quantification of spine density in neural cultures pre-treated with 50 μM KYNA (24h) and co-cultured with iMG cells. Cultures pre-treated with 50 μM KYNA showed a signifcant decrease in spine density (p=0.014). (G) Combining the pre-treatment of 50 μM KYNA with 100 μM NMDA and 250 nM D-serine reversed the effect of KYNA on spine density. Experiments in (F, G) were performed on day 25 neurons in duplicates and repeated three times. Data in all graphs (E-G) are normalized to the control group. All error bars in the figure indicate standard errors of the mean (SEM). Data was analyzed using Mann Whitney *U* tests. To exclude effects of KYNA on spine density before adding iMGs, we also performed the same experiments without adding iMGs (Figure S1), and adjusted baseline spine densities. All reported p-values are two-sided, statistical significance set to p<0.05 after adjustment for multiple testing.

### KYNA decreases spine density in human neuron-microglia co-cultures

To investigate how KYNA influences spine density, we proceeded to a co-culture system by adding human microglia-like cells to neuronal cultures pre-treated with 50 μM KYNA for 24 hours.^4^ In the co-culture system, we then observed a significant decrease in spine density (Figure 1F). To specifically address whether this observation was mediated by the effect of KYNA on NMDAR, we also performed experiments by adding 100 μM NMDA and 250 nM D-serine. This revealed that NMDA or D-serine prevented the effect exerted by KYNA on synapse density (Figure 1G).

### Spontaneous microglial engulfment of synaptic structures in forebrain organoids

To more closely mimic relevant human *in vivo* conditions, we used a human iPSC-derived dorsal forebrain organoid model. To promote the innate development of microglia within these organoids, while limiting the generation of unwanted mesoderm-derived cells, we generated iPSC-derived NPCs (from neural rosettes) and primitive macrophage progenitors (derived from yolk sac embryoid bodies) in parallel. These cells were then combined to form multi-lineage forebrain organoids^21^ by sequential addition of factors for neuronal and microglial differentiation (Figure 2A). Characterization of the forebrain organoids by single cell RNA sequencing and immunohistochemistry was performed at 120 days *in vitro* (DIV) and supported the development of mature neuronal cell types in addition to microglia (Figures 2B, 2C, 2D, S2). Employing confocal imaging, we observed the spontaneous uptake of synaptic structures in microglia (Figure 2E), substantiating a human synaptic pruning model that closely replicates *in vivo* conditions.

**Figure 2:**
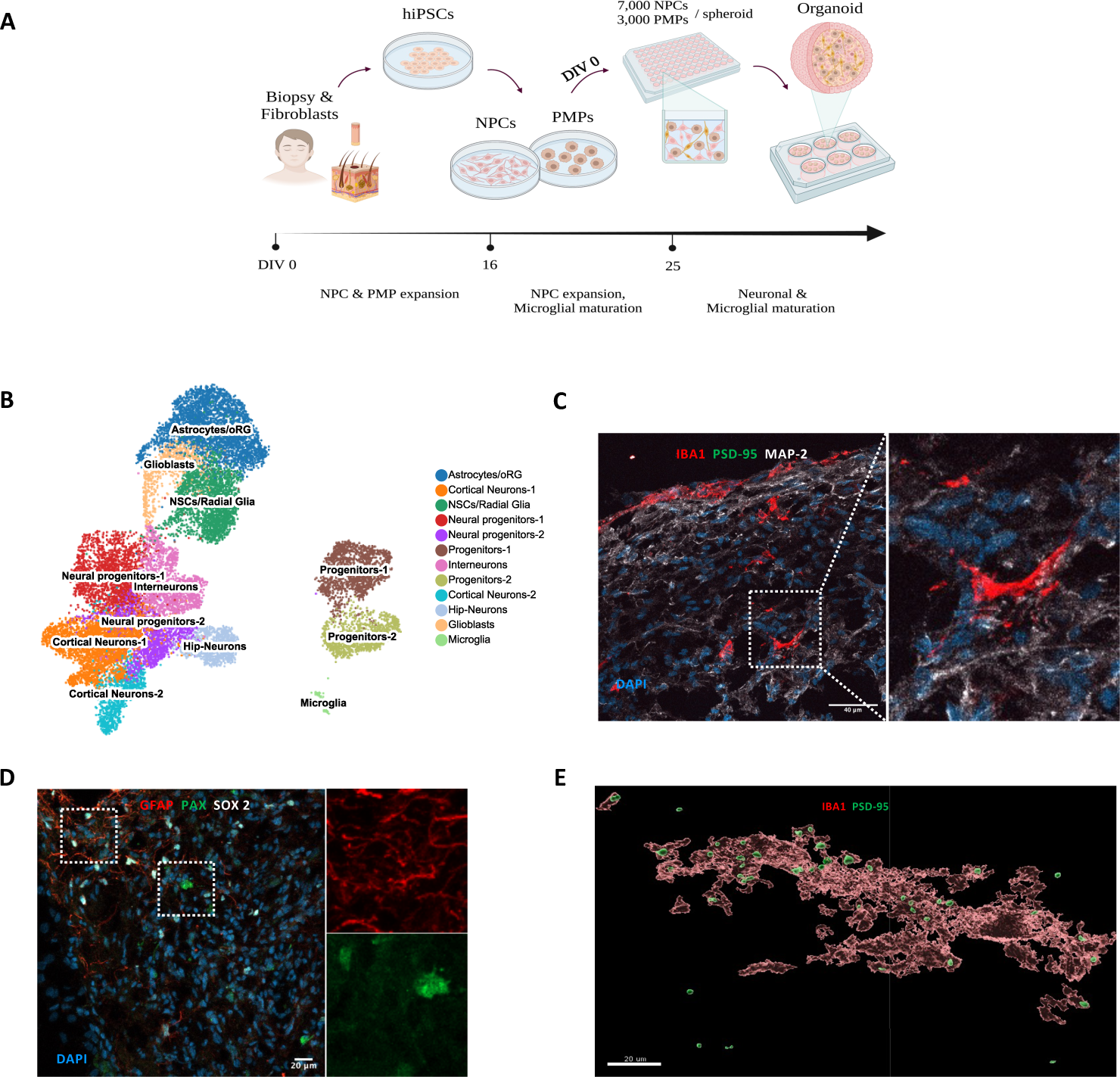
Spontaneous microglial synaptic pruning in human forebrain organoids. A) Schematic of the methods used to generate patient-derived multi-lineage dorsal forebrain organoids. B) UMAP embedding of *33,204* cells from human forebrain organoids (120 DIV) colored by 12 unique cellular clusters and confirming the presence of neuronal and glial cell types in organoids. Each dot represents a cell. C) Representative confocal images (40x) of 20-μm-thick organoid sections containing microglia (IBA1) and neurons (MAP2, PSD-95). Scale bar; 40 μm. D) Representative confocal images (20x) of 20-μm-thick organoid sections displaying astrocytic lineage cells (GFAP), NPCs (PAX) and early progenitor cells (SOX2). Scale bar; 20 μm. C) Nuclei are counterstained with 4′,6-diamidino-2-phenylindole (DAPI) in both images. E) Imaris based 3D reconstruction displaying the spontaneous overlap between post synaptic termini (PSD-95) and microglia (IBA-1) in organoids. Scale bar; 20 μm.

### Inhibiting the production of KYNA leads to decreased microglial synaptic pruning

To evaluate the effect of KYNA on synaptic pruning we generated organoids from 3 individuals with SCZ (clozapine treated) and at 163 DIV induced endogenous formation and release of KYNA by adding 1 μM of the immediate precursor kynurenine alone or in combination with 50 μM of the KAT enzyme inhibitor PF-04859989 (dose inhibiting all four KAT enzymes)^22,23^ for 48 hours. The presence of 50 μM PF-04859989 was associated with a decrease in KYNA levels (from low nanomolar concentrations) for each subject (32 %, 47 %, and 52 %, respectively: Figure 3A), confirming a human model in which we could assess how manipulation of endogenous KYNA formation influences microglial synapse pruning. In patient-derived forebrain organoids with inhibited KAT enzymes, we observed significantly longer average distances between microglia and postsynaptic structures compared to non-inhibited ones (Figure 3B), and a concomitant and significant decrease in microglial uptake of synaptic structures (Figure 3C and 3D), confirming that modulation of KYNA levels already in the nanomolar concentration range influences microglial engulfment of synaptic structures.

**Figure 3:**
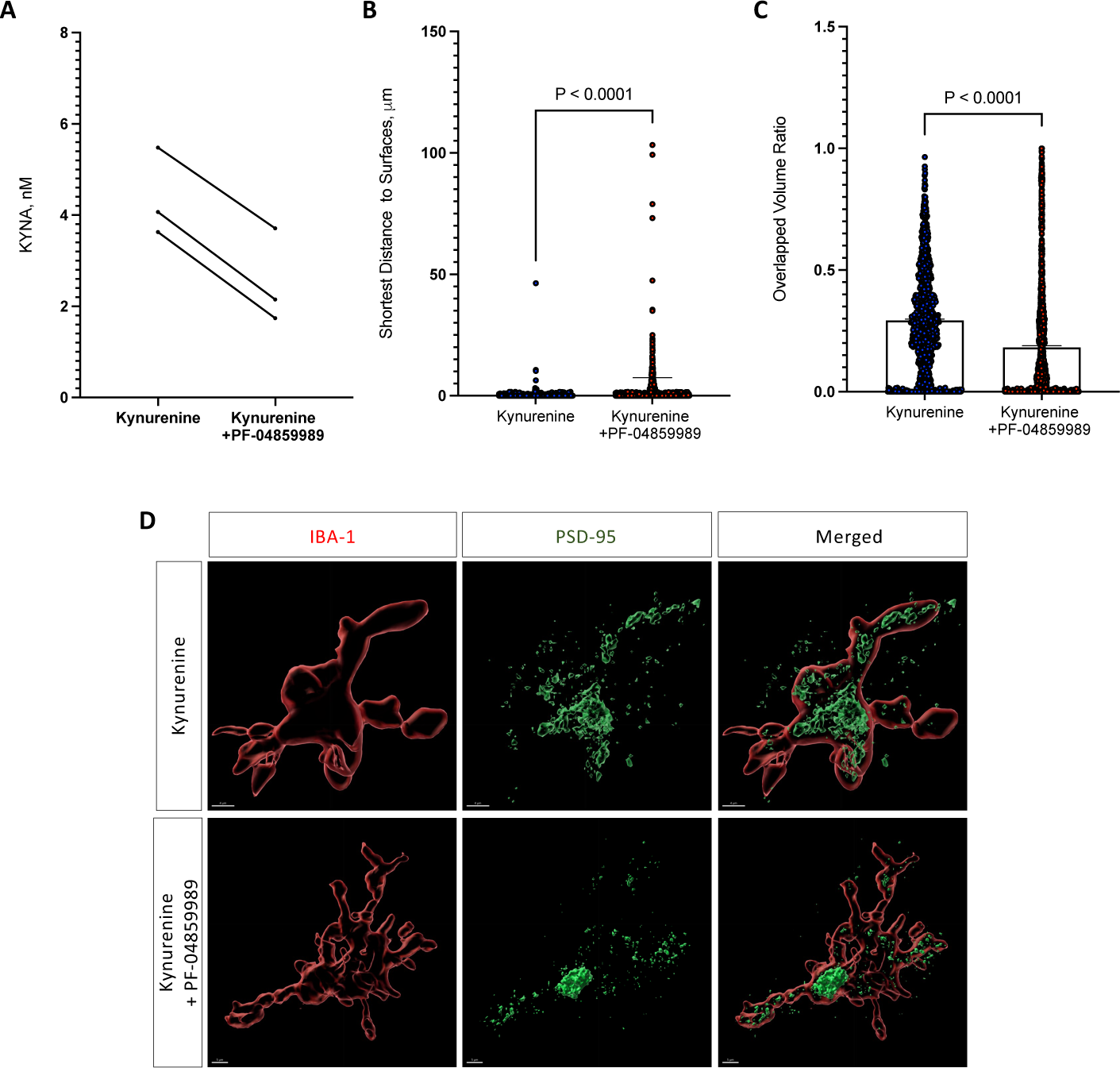
Kynurenic acid (KYNA) promotes microglia-mediated synaptic pruning in human forebrain organoids. A) Human dorsal forebrain organoids with innately developed microglia exposed to 1 μM kynurenine alone or in the presence of 50 μM PF-04859989 for 48 hours (n=3). PF-04859989 decreased the production of KYNA in organoids derived from each subject. B) After treatment with PF-04859989 (48 hours), microglia (IBA-1) displayed longer average distances to synaptic structures as indicated by the post-synaptic marker PSD-95 (p<0.0001), and C) a decreased uptake of synaptic structures (PSD-95) (p<0.0001). N=2 subjects with SCZ. (D) Representative 3D reconstructed images (Imaris) displaying the overlap between post synaptic termini (PSD-95) and microglia (IBA-1) in organoids treated with kynurenine or kynurenine and PF-04859989. Data were analyzed using Mann-Whitney *U* tests. All error bars indicate standard errors of the mean (SEM). All reported p-values in the figure are two-sided, statistical significance set to p<0.05.

### KAT co-expression networks enrich for synapse-related pathways in human prefrontal cortex tissue

Four KAT isoforms (KAT-I to -IV) are known to perform the irreversible transamination of the pivotal intermediate kynurenine to form KYNA.^24^ In contrast, kynurenine 3-monooxygenase (KMO) catalyzes the degradation of kynurenine to 3-hydroxykynurenine (3-HK) and instead promotes the production of the downstream metabolite quinolinic acid (QUIN).^24^ As KYNA is unable to cross the blood-brain barrier, endogenous brain KYNA levels are then largely reflected by KAT and KMO activity in the brain.^24^ To take advantage of this, we used large-scale brain transcriptomic data (n=224 adult frontal cortex samples; GTEx^25^ to functionally annotate each of the genes that code for different KAT isoforms (KAT-1: *CCBL1*, KAT-II: *AADAT*, KAT-III: *CCBL2*, and KAT-IV: *GOT2*) as well as *KMO*. Unsupervised co-expression network analyses were performed and ‘seeded’ on each gene to generate positively, and negatively correlated gene expression networks (Figure 4A), then expected to include the neuro-modulatory actions of endogenous KYNA in the human frontal cortex.^26^ In total, we identified 2285 *CCBL1-*positive genes and 2012 *CCBL1-*negative genes (FDR <0.05; Figure 4B), 1955 *AADAT*-positive genes and 1931 *AADAT*-negative genes (FDR <0.05; Figure 4C), 1132 *CCBL2-*positive genes and 833 *CCBL2-*negative genes (FDR <0.05; Figure 4D), while the *GOT2-*positive network contained 6513 *GOT2*-positive genes and 5650 *GOT2-*negative genes (FDR <0.05; Figure 4E), and 11950 *KMO*-positive genes and 10490 *KMO*-negative genes (FDR<0.05; Figure 4F and Table S1). *CCBL1* and *GOT2* positive genes, as well as *AADAT* negative genes, also included *GRIN1* (coding for the glycine site carrying subunit of NMDAR) and displayed significant correlations with expression of these KAT genes after FDR adjustment (Table S2). In addition, *GOT2* and *KMO* expression positively correlated with *CHRNA7* expression (coding for the subunit of α7nAchR). In line with previous findings, our findings support the hypothesis that the antagonistic properties of KYNA on the glycine site of NMDAR^27^ and α7nAchR^28^ are relevant in a human *in vivo* context.

**Figure 4.**
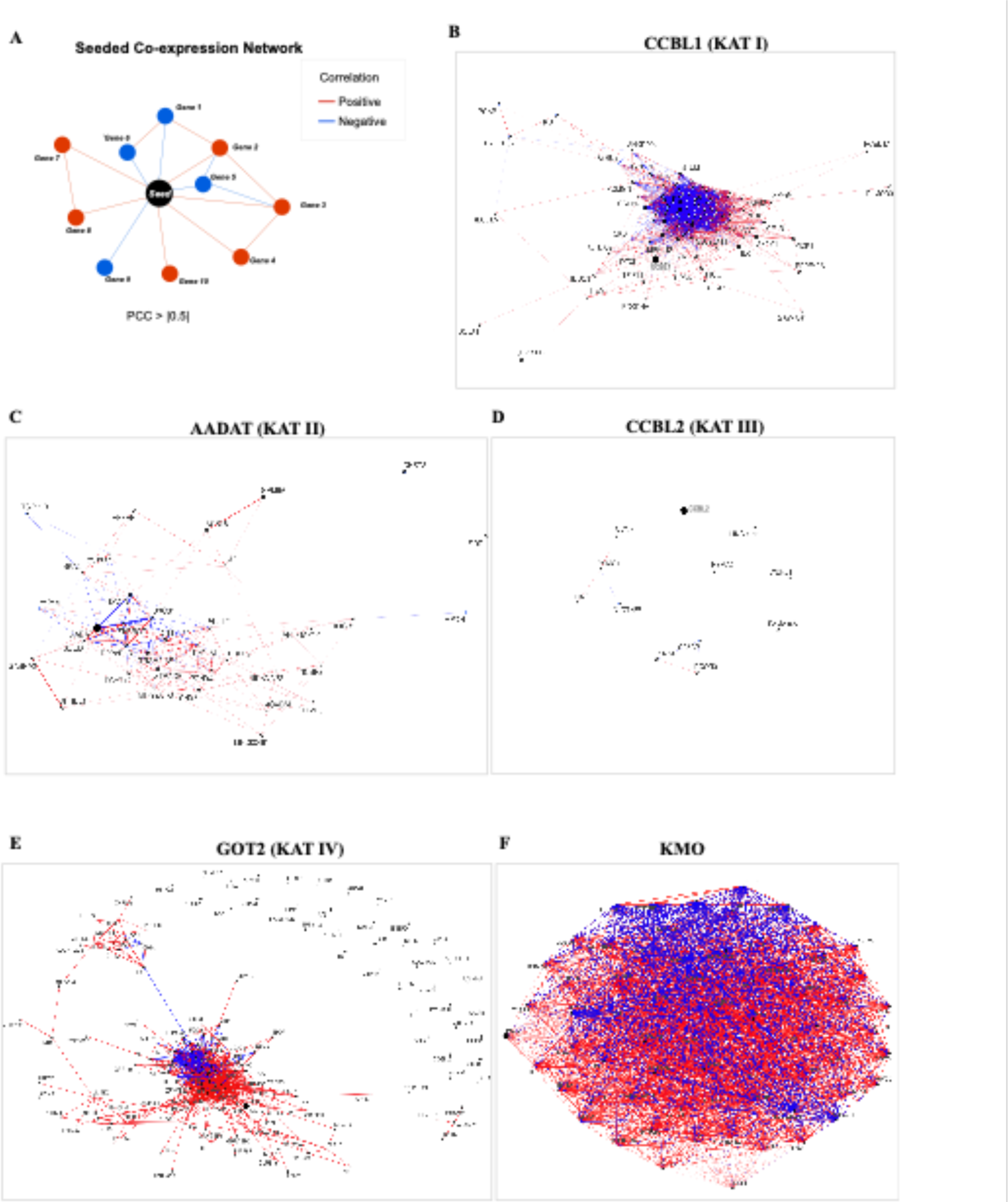
Overview of unbiased seeded frontal cortex co-expression networks. (A) Schematic of the constructed seeded co-expression networks. (B) *CCBL1* (KAT I), (C) *AADAT* (KAT II), (D) *CCBL2* (KAT III), (E) *GOT2* (KAT IV), (F) *KMO*, used as the ‘seed gene’ (black dot), in frontal cortical samples of neurotypic individuals (*n=224*).^25^ Red lines represent the positive network while blue lines represent the negative network of the respective seed gene. Edges denote the gene-gene co-expression. The Pearson’s correlation coefficient (PCC) threshold was set to 0.5. Transcriptome-wide significance was calculated at FDR-adjusted p-values<0.05 (*20,004* detected genes).

Next, we investigated the identified co-expressed networks for the enrichment of functional pathways as represented by unbiased global gene modules computed within the same transcriptomic dataset. Both *CCBL1, GOT2,* and *KMO* positive networks significantly enriched for modules with synapse-related pathways, whereas the *AADAT* negative network displayed enrichment for the same modules (Figure 5A, 5B and Table S2). Except for *AADAT*, all other KAT negative networks were also enriched for immune-related pathways (Figure 5A and Table S3), while *GOT2*-positive genes exhibited substantial enrichment for modules involving pathways that regulate synapse assembly (Figure 5A and 5B). This largely aligned with the developmental expression trajectories (single-cell fetal dataset: N=48 prefrontal cortex samples,^29^ and single-cell postnatal dataset: N=16 prefrontal cortex samples,^30^ displaying a dominant *GOT2* expression during the fetal period, while in late adolescence (*CCBL1/2*) and early adulthood (*AADAT*), i.e., the time period largely overlapping with extensive synaptic pruning in the prefrontal cortex,^31^ and the typical age of onset for SCZ,^32^ *GOT2* expression decreased while the expression of the other KAT genes simultaneously increased (see Figure S3 for overall expression trajectories of the KAT genes and *KMO* while expression profiles in single-cell types can be found in Figure S4). Furthermore, in the fetal dataset, we could confirm significant enrichment of excitatory neurons in the positive networks of *CCBL1* and *GOT2* (Figure S5).

**Figure 5.**
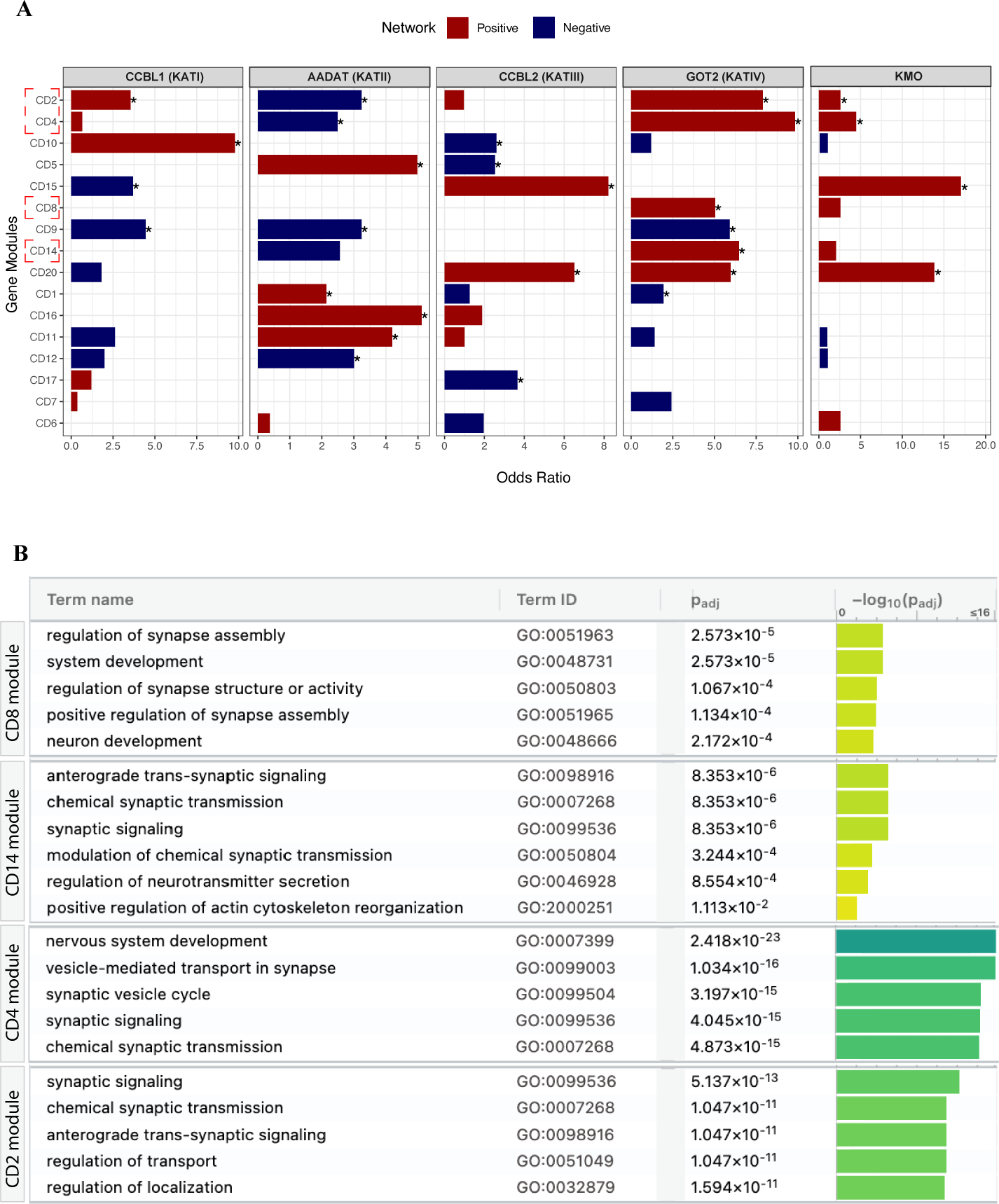
Enrichment of functional pathways in seeded frontal cortex co-expression networks. (A) Bar plots displaying the enrichment of the global WGCNA gene modules (CD) (y-axis) in the top correlated networks of individual *KAT*-enzymes and *KMO* (top bar). X-axis denotes the odds ratio calculated by Fisher’s exact test. Asterisks denote significant at FDR-adjusted p-values<0.05. Red bars indicate the positive network and blue bars indicate the negative networks for the respective genes (top bar). Module enrichment results are provided in Table S2. Red brackets on y-axis highlight modules related to synapse mechanisms. (B) Over-representation analysis biological pathways (Gene Ontology: BP; Term name and ID) in selected gene modules, implicating synapse-related pathways, that are significantly enriched in KAT co-expression networks (from figure 5A) using hypergeometric tests. Enrichment p-values were corrected for multiple testing using the FDR method (P_adj_) and plotted as −log(P_adj_) on the y-axis of the bar plot. Module names are indicated on the left vertical bar.

### KAT co-expression networks enrich for schizophrenia risk variants

Despite repeated meta-analyses implicating elevated brain levels of KYNA in SCZ,^17,33,34^ a link between genetic risk variants, i.e., the main risk factor for SCZ, and brain KYNA levels remain to be proven. To address this, we integrated the unbiased KAT enzyme co-expression networks with genetic risk genes from two large-scale GWAS analyses,^35–37^ and one trio-based exome sequencing study of SCZ.^38^ To differentiate unique and cross-disorder risk variants, we also used a SNP-based autism (ASD) risk gene set.^38^ *CCBL1-* positive networks displayed a strong and specific enrichment for SCZ risk (Figure 6A; all SCZ risk gene sets but not the autism risk gene set) while *AADAT*-positive networks displayed a significant enrichment for one of the GWAS-based risk gene set (Figure 6B). *CCBL2* networks displayed no significant enrichment for any of the risk gene sets (Figure 6C), while *GOT2*-positive networks displayed enrichment for both SCZ risk genes (GWAS) and ASD (Figure 6D). Additionally, *KMO-* negative networks exhibited enrichment for SCZ risk genes (Bonferroni-corrected significance; Figure 6E). Comparing the list of enriched risk genes (focusing on *CCBL1* and *AADAT* positive networks) with genes driving the enrichment for functional synapse-related pathways, we observed no significant overlap. Instead, the enriched risk genes contained genes such as *IL17RB* (interleukin 17 receptor B), *RFTN2* (Raftlin Family Member 2), and *AMBRA1* (autophagy/ beclin-1 regulator 1 gene), all capable of increasing the expression of cytokines that can induce the formation of KYNA upstream of KATs.^24,39,40^ This suggests that elevated KYNA levels in SCZ are partly due to common genetic risk variants, and that the enrichment for synapse-related pathways is not a direct consequence of co-expression with synapse-related SCZ risk genes but rather a consequence of elevated KYNA levels.

**Figure 6.**
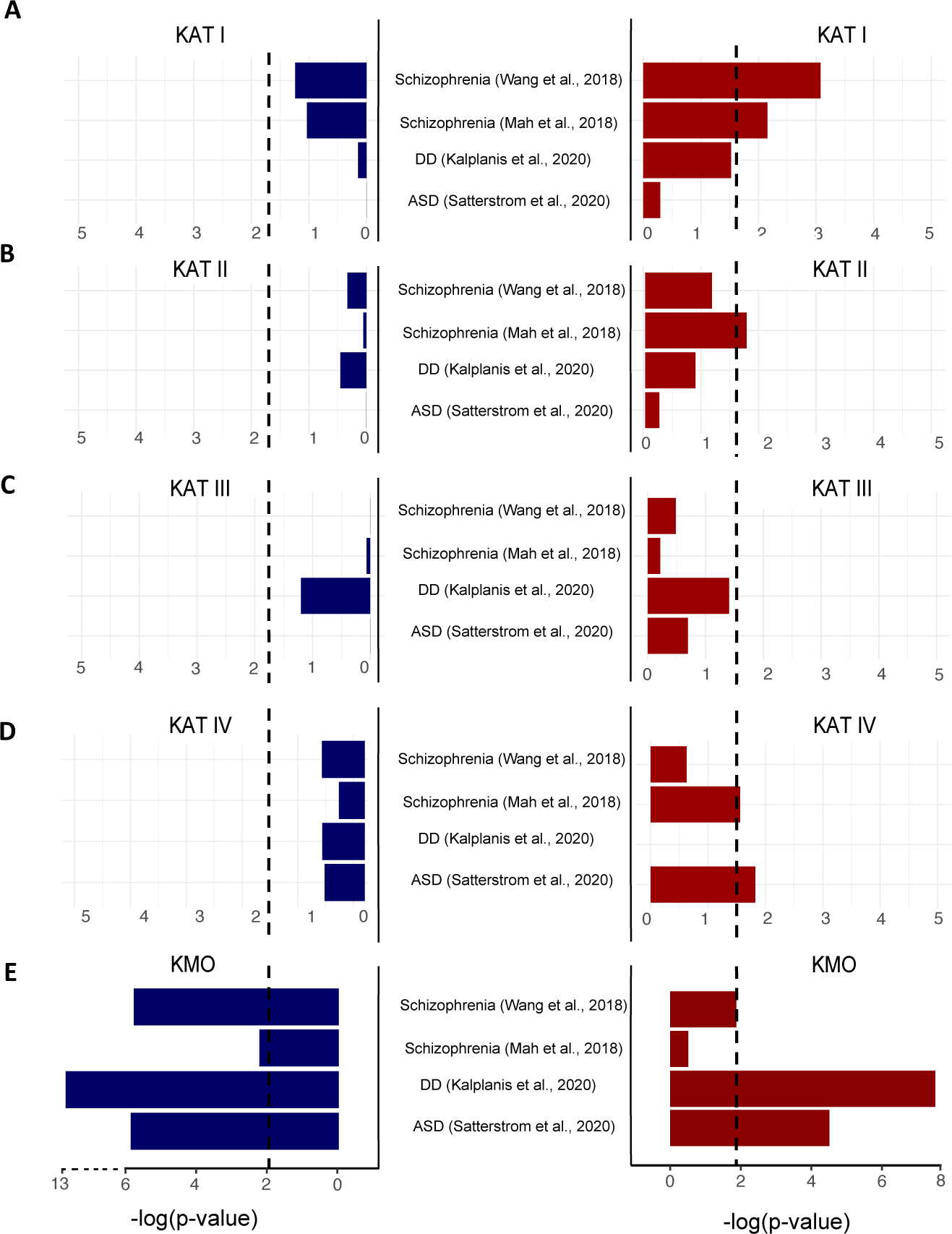
Enrichment of disease-specific risk genes in the seeded frontal cortex co-expression networks. Including two schizophrenia (SCZ) risk gene sets,^35–37^ an autism spectrum disorder (ASD),^38^ and a developmental disorder (DD)^66^ risk gene set in computed co-expression networks of (A) KAT I, (B) KAT II, (C) KAT III, (D) KAT IV, and (E) KMO. Enrichment p-values (fishers exact test) are plotted as −log_10_(p-values) on the y-axis. Dotted line represents Bonferroni threshold (*n=4*) for multiple tests adjustment. Risk gene sets obtained from published studies are shown on x-axis. Red bars (right) show enrichment of disorder risk in *KAT*-positive networks and blue bars (left) show the enrichment of disorder risk in *KAT*-negative co-expression networks.

## Discussion

This study defines a role of KYNA in promoting microglial synapse removal by inhibiting glutamatergic activity. We first used a 2D model to show that excitatory neurons treated with KYNA displayed a reduction in spine density when co-cultured with microglia and that this effect can be reversed by adding NMDA. We then used a 3D approach and inhibited the production of KYNA in forebrain organoids and observed a decrease in microglial uptake of synaptic structures. Utilizing human large-scale forebrain transcriptomic datasets, we confirmed that the expression of KYNA-producing enzymes correlated with the expression of NMDA and α7nAchR subunits, while the seeded co-expression networks enriched for synapse-related pathways and excitatory neurons. To address risk factors causing elevated KYNA levels in SCZ, we then took advantage of seeded co-expression networks and showed enrichment for genetic SCZ risk based on common risk variance.

KYNA concentrations in the mammalian brain typically reside in the nanomolar range.^41^ Although local concentrations at the synapse can reasonably be expected to be substantially higher, it is unlikely that the millimolar concentrations needed to inhibit the glutamate site of NMDAR are reached.^10,42^ Thus, we here first used micromolar concentrations of KYNA (2D), to reach concentrations at the synapses for which the antagonistic properties of KYNA on the glycine co-agonist site of the NMDAR and on α7nAchR are well-established (but excluding effects on the glutamate site of NMDAR).^42^ In the organoid model, we then took advantage of the innate production of KYNA in these self-organizing 3D cultures and stimulated the KAT enzymes by further increasing kynurenine levels. This resulted in low nanomolar concentrations of KYNA in the organoids, i.e., mimicking average KYNA concentrations observed in the human brain. Our large-scale analyses of human postmortem frontal cortex tissue then also confirmed strong correlations between expression of KAT enzymes and genes coding for subunits of the NMDAR as well as *CHRNA7*, further supporting that the effects of KYNA in our model systems are relevant also in a human *in vivo* context.

Repeated observational studies have reported elevated KYNA levels in the CNS of patients with SCZ,^17^ while animal studies have revealed that KYNA can induce cognitive deficits such as spatial learning and memory impairments that can be observed in SCZ,^11,43,44^ i.e., phenotypes that have been linked to excessive microglial synaptic pruning in animals.^45,46^ The discovery that KYNA promotes microglia-mediated synaptic pruning then defines a precise and plausible molecular mechanism linking elevated KYNA levels to the observed cognitive deficits in patients with SCZ. It is important to note that a C4A induced sensitization of the microglial synapse recognition system in SCZ to some extent overlaps with this activity-driven mechanism mediated by KYNA as interleukin (IL)-1β elevated in SCZ^47^ both stimulates the expression of *C4A*^48^ and the production of KYNA (by inducing expression of tryptophan-2,3-dioxygenase upstream of the KATs).^49^ In line with an integrated mechanism, C4A negative co-expression networks, as the KAT-positive networks, also enrich for synaptic pathways that involve SCZ risk variants.^50^ As it has been challenging to directly target the complement system in SCZ, expanding the search area to synapse-related mechanisms will introduce new more accessible drug targets.

Several isozyme-specific inhibitors of KATs (mainly focusing on KAT-II) have been shown to decrease brain KYNA levels and are under development for clinical application.^23,51–53^ Increased activity of both KAT-I and KAT-II, as well as decreased activity of KMO, have also been linked to elevated KYNA levels under pathophysiological brain conditions,^54^ while increased *CCBL1* and *AADAT* expression, as well as decreased *KMO* expression, have been observed in postmortem brain tissue obtained from patients with SCZ.^54,55^ Our current findings suggest that KAT inhibition as a treatment strategy for SCZ should take place in the early phases of the disorder when a substantial loss of synapses can be observed.

## Material and methods

### Ethical statement

All skin biopsies and blood samples were obtained with written informed consent. The study was approved by the Regional Ethics Committee of Stockholm (Sweden) and the Ethics Committee of the Helsinki University Hospital District. The study was carried out in accordance with the latest Helsinki guidelines.

### Generation of iPSCs

iPSCs were generated from dermal fibroblasts either using mRNA or Sendai virus-based reprogramming.^56^ Skin biopsies were collected from patients with SCZ (age range: 28-47, 2 females and 2 males).

### Generation of *NGN2* NPCs

iPSC colonies were stabilized under xeno-free culture conditions, plated on biolaminin 521 LN-coated (BioLamina) plates, and expanded in NutriStem XF medium (Biological Industries) to passage 3 or more. TRA-1-60^+^ iPSCs were isolated from cultures using anti-TRA-1–60 MicroBeads (Miltenyi Biotec) and LS columns (Milteny Biotec), according to the manufacturer’s instructions. Obtained iPSCs were characterized by octamer-binding transcription factor 4 (POU domain, class 5, transcription factor 1) and TRA-1–60 immunostaining.

Inducible neurogenin 2 (NGN2)-expressing expandable neural progenitor cells (NPCs) were obtained using TALEN-based plasmids as described earlier.^4^ Briefly, the doxycycline-inducible *NGN2* AAVS1 knock-in plasmid, based on an AAVS1 SA-2A-puro-pA donor (plasmid no. 22075; Addgene), was generated by replacing the puromycin resistance gene with the neomycin resistance gene and cloning a cassette containing the Tet-On 3G promoter driving the human *NGN2* complementary DNA followed by P2A, a zeocin resistance gene, the BGH polyA, CMV early enhancer/chicken β-actin (CAG) promoter driving Tet-On 3G and BGH polyA. The AAVS1-targeting TALENs are based on hAAVS1 1R TALEN and hAAVS1 1L TALEN (plasmid nos. 35432 and 35431, respectively; Addgene) and were generated by golden gate assembly. Plasmid maps can be provided upon request. NPCs were collected with Accutase (ThermoFisher Scientific) and counted. For each transfection, 4−10^6^ cells were pelleted at 300xg and then resuspended in 100 μL prewarmed nucleofector solution (Human Stem Cell Kit 1, Lonza, catalog no. VPH-5012); 4 μg of NGN2 plasmid and 1.5 μg each of 1R and 1L plasmids were added directly to the resuspended cells, or negative control without plasmid, followed by nucleofection (Amaxa Nucleofector II; Lonza) according to the manufacturer’s protocol using program B-016. After nucleofection, cells were plated onto Geltrex-coated 6-well plates in neural expansion medium (50% NBM, 50% advanced DMEM/F-12 with 1x NIS). Stable lines were expanded by selection with the addition of 125 μl mL^−1^ G418 (Geneticin; Thermo Fisher Scientific) over 8–10 days or until the negative transfection control cells completely died.

### Generation of excitatory neurons

*NGN2* induced neurons were derived from NPCs stably expressing *NGN2* as described earlier.^4^ Briefly, NPCs were plated as single cells on poly-L-ornithine/laminin coated 24-well polypropylene plates at 2.5−10^5^ cells/500 μL in 50% neurobasal medium (NBM; ThermoFisher Scientific) and 50% DMEM/F12 (ThermoFisher Scientific) with 1x neural induction supplement (NIS; ThermoFisher Scientific). One day after seeding, the medium was replaced with neuronal differentiation medium (50% NBM and 50% DMEM/F12 supplemented with 1x MEM Non-essential Amino Acids (ThermoFisher Scientific), 1x N-2 supplement (ThermoFisher Scientific), 10 ng mL^−1^ brain-derived neurotrophic factor (BDNF; Peprotech) and 10 ng mL^−1^ recombinant human NT-3 (PeproTech)). Two days after seeding, the medium was changed to fresh differentiation medium enriched with 1x B-27 (ThermoFisher Scientific) and 2 μg mL^−1^ doxycycline (Sigma-Aldrich). This medium was maintained for the entire differentiation process, unless noted. On days 3-6, cells were incubated with the same medium with the addition of 5 μg mL^−1^ zeocin (ThermoFisher Scientific). For neuronal maturation, eleven days after seeding, astrocyte conditioned medium (ScienCell Research Laboratories) was added to the culture. Synaptic pruning assay was then performed at day 25.

### Generation of induced microglia-like (iMG) cells

Whole blood was collected into vacutainer cell preparation tubes containing sodium citrate as an anticoagulant (Becton, Dickinson and Company) and processed as per the manufacturer’s instructions. Peripheral blood mononuclear cells (PBMC) were isolated, washed twice with PBS by centrifugation and suspended in heat-inactivated fetal bovine serum (FBS; Sigma-Aldrich) containing 10% DMSO (Sigma-Aldrich). The cell suspension was then divided into aliquots, transferred into cryovials and cryopreserved by using a liquid nitrogen cooled freezer.

Generation of induced microglia-like cells (iMG) from PBMCs was carried out using methods previously described,^57^ with minor modifications. Briefly, cryopreserved PBMC samples were transferred from liquid nitrogen freezer to a 37 °C water bath. Once the cell suspension had been thawed, it was gently pipetted into 10 mL of pre warmed complete RPMI medium consisting of basal RPMI-1640 supplemented with 10% heat-inactivated FBS and 1% penicillin-streptomycin (P/S; ThermoFisher Scientific). Following centrifugation (300x*g* for 5 min at room temperature (RT)), the supernatant was discarded, and the cell pellet was resuspended in the appropriate volume of complete medium.

Isolated PBMCs were cultured at a density of 5−10^5^ cells/1 mL complete medium on 24-well plates coated with Geltrex (Thermo Fisher Scientific). After 24 hours of incubation, the media was replaced with RPMI-1640 supplemented with 1x glutamax (Life Technologies), 1% P/S, 0.1 μg mL^−1^ of interleukin (IL)-34 (Peprotech) and 0.01 μg mL^−1^of granulocyte macrophage colony-stimulating factor (GM-CSF; Peprotech). After 7 days, fresh media was added. At day 11, iMGs were used for co-culture experiments.

### Characterization of excitatory neurons

*NGN2* induced neurons were cultured on glass microscope slides coated with poly-L-ornithine/laminin and fixed in 4% paraformaldehyde in PBS for 15 min. Following washing with PBS and permeabilization in PBS containing 0.1% Triton-X100 for 5 min, cells were blocked with blocking solution (PBS containing 5% normal goat serum) for 1 hour at RT. Primary antibodies against MAP-2 (1:500, Abcam), Beta III Tubulin (1:2000, Promega), and PSD-95 (1:100, Abcam) were diluted in blocking solution and added to the cells. After overnight incubation 4°C, cells were then washed with PBS and incubated with respective Alexa Fluor-conjugated secondary antibodies (1:500) for 1 hour at RT. DAPI (1:5000) was added as a nuclei/DNA marker for 10 min at RT. Images were obtained using a confocal microscope system (Zeiss, LSM900-Airy2).

### Calcium Imaging

For the detection of intracellular calcium levels, *NGN2* induced neurons were cultured on black Corning imaging 96-well-plates (Sigma-Aldrich) and maintained as per culture conditions. Neurons at 22 days after induction were treated with 25 or 50 a KYNA for 24 hours. The calcium indicator was prepared by mixing 50 μg Fluo-3/AM (Invitrogen) with 50 uL DMSO and vortexed. 50 μL of Pluronic™ F-127 (Invitrogen) was added and vortexed. 5 uL of this composition was mixed with 600 uL neuronal differentiation medium. Old media was removed from the cultures and the mixture with the calcium indicator was added to each well and incubated for 15 min at 37 °C with 5% CO_2_. Following incubation, cells were washed and assay buffer (20 mM HEPES, 2 mM CaCl_2_, 119 mM NaCl, 11 mM Glucose, and 2.5 uM KCl with autoclaved water in a final volume of 50 mL, pH 7.4) was placed to each well. Intracellular calcium level detection was performed on Clariostar (BMG Labtech) (excitation 483 nm and emission 530 nm).

### Co-cultures and quantification of spine density

After 25 days of induction as described, neural cultures were treated with 50 μM KYNA alone or in the presence of 100 μM NMDA and 250 nM d-serine for 24 hours and washed with PBS prior to co-culture experiments. Mature iMG cells were collected from plates using Accutase, pelleted and counted; 3−10^4^ iMG cells were added to each neural culture well. After 24 hours of interaction, co-cultures were washed with PBS and fixed with 4% paraformaldehyde (PFA) for 15 min and stained with Alexa Fluor 488 phalloidin (Thermo Fisher Scientific) for 40 min at RT. Images from neural-iMG co-cultures were acquired using an ImageXpress Micro automated microscope (Molecular Devices) and analyzed using ImageJ software (version 1.52). Sixteen images per well at 20x magnification were taken, using FITC channel (excitation 482/35 nm and emission 536/40) to capture phalloidin 488 fluorescence signal. Acquired TIFF files were processed using a customized macro in imageJ.

### Generation of forebrain organoids

Forbrain organoids were generated using the protocol from Xu et al.,^21^ with some modifications. Briefly, for generation of PMPs, 10^4^ iPSCs were seeded per well on ultra-low attachment V bottom 96-well plates in mTeSR medium supplemented with rock-inhibitor (Sigma-Aldrich). Twenty-four hours after seeding, embryoid bodies (EBs) were formed and media was carefully changed to ‘EB media’ containing mTeSR supplemented with 1 mM rock-inhibitor, 50 ng mL^−1^ BMP-4 (Peprotech), 20 ng mL^−1^ SCF (Peprotech) and 50 ng mL^−1^ VEGF (Peprotech). EBs were carefully observed over the next four days and media were changed every day using freshly prepared EB media. After four days, using a cut 200 ul pipette tip, EBs were transferred onto a tissue culture treated 6-well plate (20-25 EBs per well) into ‘pMACpre media’ containing X-VIVO15 [with recombinant transferrin, w/o gentamycin and phenol red] with 100 ng mL^−1^ M-CSF, 25 ng mL^−1^ IL-3, 1x GlutaMAX, 1x Pen-Strep, 0.055 μM BME). Media was changed weekly using pMACpre differentiation media for four weeks. After about three weeks, pMACpre cells started appearing in suspension. pMACpre cells were further harvested and combined with pNPCs to form organoids as described in the section below. As a quality control measure, to test if the cells could differentiate into microglia, PMPs were harvested from the supernatant and plated on geltrex coated 24-well plates in ‘pMGL medium’ containing Advanced DMEM/F-12 with N2, 100 ng mL^−1^ IL-34, 10 ng mL^−1^ GM-CSF, 1x Glutamax, 1x Pen-Strep and 0.055 μM BME. After two weeks, microglial morphology was observed. PMPs cannot be frozen so cells were used fresh for every batch of forebrain organoids.

For generation of iPSC-derived NPC we first seeded Tra-1-60 purified iPSCs on ultra-low attachment V bottom 96-well plates (10^4^ cells per well). EBs were formed within 24 hours and the media was carefully changed using freshly prepared media containing DMEM/F12 supplemented with 1x N2, dorsomorphin (1 μM) and SB431542 (5 μM). Media was changed every other day for one week. EBs were then carefully transferred (using a cut 200 μL pipette tip) onto growth factor reduced matrigel coated 6-well plate (6 EBs per well) and containing media composed of DMEM/F12 supplemented with 1x N2, and 1 mg mL^−1^ laminin 521. Media (DMEM/F12 supplemented with 1x N2) was carefully changed every other day. After 7 days, neural rosettes were observed and harvested manually from surrounding cells. Rosettes were plated on growth factor reduced matrigel coated 6 well plates in ‘pNPC medium’ composed of a 1:1 mixture of Neurobasal (ThermoFisher Scientific) and DMEM/F12, supplemented with 1x N2, 1x B27-RA (ThermoFisher Scientific), 20 ng mL^−1^ FGF2 (Peprotech), 3 μM CHIR99021 (Biogems), 10 ng mL^−1^ human leukemia inhibitory factor (hLIF, Millipore), 2 μM SB431542, and 10 μM rock-inhibitor. pNPCs were passaged twice using Accutase. In order to purify the NPCs, PSA-NCAM positive cells were obtained using magnetic bead separation kit from Miltenyi Biotech and plated on geltrex coated 6-well plates at a concentration of 500,000 cells per well. When pNPCs were confluent, they were again magnetically sorted to obtain CD271-CD133+ cells. These purified pNPCs were used for generation of COs.

PMPs and NPCs were then combined to generate forebrain containing microglia. PMPs were harvested from the supernatant using reversible 37-micron strainers. Cells were centrifuged at 300xg for 5 min and resuspended in 1 mL of ‘proliferation media’ which was composed of a 1:1 mixture of ‘pMACpre media’ and ‘pNPC media’. At the same time, NPCs were harvested using Accutase, resuspended in 1 mL of proliferation media and counted. Seven thousand pNPCs and 3000 PMPs cells were plated per well of an ultra-low attachment 96-well plate. Each well produced one organoid. The plate was centrifuged at 150xg for 5 min and incubated at 37 °C CO2 incubator. This is considered as day 21 in the organoid protocol. Proliferation media was changed very carefully every day until day 24. Using a cut 200 μl pipette tip, each organoid was then carefully transferred to an ultra-low attachment 24-well plate; one organoid per well. Proliferation media was changed every day until day 27 when the organoids were carefully transferred to an ultra-low attachment 6-well plate (4 organoids per well) containing 3 mL of proliferation media; and placed on a shaker at 90 rpm in the incubator. Proliferation media was changed very carefully, every day until day 35 when the media was changed to ‘maturation media’ containing 50% neurobasal medium, 25% DMEM/F12, 25% BrainPhys medium supplemented with 20 ng mL^−1^ BDNF, 20 ng mL^−1^ GDNF, 1 mM dibutyryl cyclic AMP, 10 ng mL^−1^ IL-34 and 1 ng mL^−1^ GMCSF. Maturation media was changed every other day until day 163.

### Characterization and quantification of synaptic pruning in forebrain organoids

Organoids were fixed in 4% PFA at 4°C for 45 min, washed twice for 5 min with PBS at RT and transferred to 30% (w/vol) sucrose solution for overnight incubation at 4°C. Sucrose solution was then removed and organoids were transferred to tissue base moulds and embedded in optimal cutting temperature (OCT) compound on dry ice to protect against freezing artifacts. Organoid blocks were stored at −80°C until used for cryosectioning. For immunohistochemistry, 20-μm-thick sections of frozen organoid tissue were prepared using a Leica cryostat. Briefly, organoid blocks were removed from −80°C and allowed to warm in the cryostat chamber to sectioning temperature for 30 min. The sectioning temperature of the blade and the chamber was optimized to −20°C. Multiple serial sections were collected in a total of 10 slides/organoid, each one containing at least one section from the front, the middle and the back of the tissue, allowing us to explore the localisation of markers across different regions of the organoids.

Cryosections of 20-μm thickness were then incubated in a blocking solution (0.3% triton-X100 and 3% BSA in PBS) for 60 min. Sections were washed with PBS and incubated with primary antibodies (diluted in 0.3% triton-X100 and 3% BSA in PBS) overnight at 4°C; anti-IBA1 (1:100, Wako), anti-MAP2 (1:500, Abcam) and anti-PSD-95 (1:100, Abcam). Sections were washed in PBS (3 times) and incubated with secondary antibodies (diluted in in 0.3% triton-X100 and 3% BSA in PBS) for 2 hours at RT in the dark. Finally, the sections were washed in PBS and incubated in DAPI (1:500) for 5 min before mounting with the immunofluorescence mounting media (DAKO).

Images were captured (8-bit) from a Zeiss LSM900 confocal microscope with ZEN software (Zeiss). For quantification of microglial engulfment of PSD-95 positive post-synaptic termini, tiled z-stacked images (25 um range, step-size 1.5 um) were acquired with 40x magnification (C-Apochromat 40x/1.20 W) for the whole organoid section. Maximum intensity projections and stitched images were generated post acquisition with the ZEN software and subsequently analyzed with Imaris software (Oxford Instruments). In Imaris, three-dimensional surfaces were reconstructed and the overlapping volume ratio and the shortest distance to surfaces between PSD-95 positive proteins and IBA-1 positive proteins were measured.

### Ultra-performance liquid chromatography tandem mass-spectrometry (UPLC-MS/MS)

Thirty microliters of cell medium as well as quality control samples were transferred to a 96-well plate and mixed with 30 μL of internal standards (IS) at 0.5 μM in 10% ammonium hydroxide (LC-MS grade) solution for 15 seconds (sec) covered with plate seal (RAPID Slit Seal, BioChromato, Japan). 60 μL of 200 mM ZnSO4 (5 °C) was added and mixed for 15 sec before 30 μL of methanol (5 °C) (UPLC-MS grade) was added and mixed for 15 sec. The mixture was then centrifuged for 10 min at 2841xg at RT. Thirty microliters of the supernatant were mixed with 30 μL of 5% formic acid in LC-MS Certified Clear Glass Vials (12 × 32 mm, Waters, product no. 186005662CV). The clear mixture was transferred to an autosampler (set to 5 °C) with 1.5 μL injection volume into the UPLC-MS/MS system.

The UPLC-MS/MS system: used was a Xevo TQ-XS triple-quadrupole mass spectrometer (Waters, Manchester, UK) equipped with a Z-spray electrospray interface and a Waters Acquity UPLC I-Class FTN system (Waters, MA, USA). Full description of the UPLC-MS/MS method has been published previously.^58^ In brief, the MS was operated in electrospray-positive multiple reaction monitoring (MRM) mode with a source temperature of 150 °C, capillary voltage of +3.0 kV, desolvation temperature 650 °C, desolvation gas flow rate 1000 l/h and detector gain 1. The UPLC Acquity HSS T3 2.1 × 150 mm, 1.8 μm column (Waters, Product Number [PN]: 186,003,540) was set to a temperature of 50 °C. Mobile phases used were A: 0.6% formic acid in water and B: 0.6% formic acid in methanol (UPLC grade). An isolator column (Waters, 2.1 × 50 mm column, PN: 186,004,476) was used to retain contaminants from the mobile phase. The flow rate was set at 0.3 ml/min and the run time for each sample was 13.0 min. The *m/z* for the MRM transitions of KYNA was 190 > 116 and for internal standard KYNA-*d*_5_, 195 > 121 and the retention time was 7.85 and 7.81 minutes respectively. The lowest level of quantification for KYNA was 1 nM and the coefficients of variance (%CV) for quality controls (n=4) within the assay (intra-assay), were less than 5%.

### Single-cell transcriptomics data processing of forebrain organoids

3D brain organoids were characterized using droplet-based single-cell RNA/ATAC sequencing on the chromium platform (10X Genomics). Briefly, organoids derived from 3 individuals were pooled (*n=6* organoids/individual) and dissociated using cold mechanical dissociation into single-cell suspensions. Nuclei were isolated as described in CG000375 _NucleiIsolationComplexSample_RevC (10X Genomics) and subjected to Iodixanol gradient centrifugation (Opti prep) to get rid of the cellular debris. High-quality nuclei suspensions were then loaded into to 3 separate wells on the chromium chip and processed according to CG000338_ChromiumNextGEM_Multiome Guide_RevE (10X Genomics), to obtain gene expression libraries. The barcoded libraries were pooled and sequenced on NovaSeq 6000 S4v1.5 (Illumina). Raw sequencing data was processed using Cellranger software v2.1 (10x Genomics) to obtain count matrices and merged using the *Aggr* function. Quality control metrics (percentage of mitochondrial transcripts < 10%; 1000 < number of transcripts < 30000, number of expressed genes > 200) were applied to the merged gene expression count matrix to filter out bad-quality nuclei and empty droplets. Transcriptomic doublets were identified and filtered out using the DoubletFinder tool. Filtered counts were normalized using the SCTransform method implemented in Seurat.^59^ Confounding variables such as mitochondrial percentage and differences in cell cycle scores were regressed out. The data was imported to Python for visualization. Batch effects introduced by cell line of origin were corrected using Batch balanced KNN data integration implemented in scanpy.^60^ Dimensionality reduction was performed using UMAP embeddings followed by nearest-neighbor graph construction and unsupervised clustering was performed using the Louvain algorithm. Cluster annotations were done using a combined approach of differential expression gene (DEG) analysis for marker genes specific to clusters, and reference mapping to primary human brain datasets. DE testing was computed using the MAST test (Adjusted p-value, based on Bonferroni correction using all genes in the dataset < 0.05). Reference mapping was done using whole transcriptome correlations clustifyr package^61^(Pearson’s correlation).

Reference single-cell RNA-seq datasets were downloaded from Nowakowski et al.,^29^ (Fetal; gestational weeks 9-39), Velmeshev et al.,^30^ (Human postnatal including adolescence; 2-24 years old. Count matrices and metadata for each dataset were processed and gene expression was visualized using Seurat R package. Cluster annotations provided by authors were validated and utilized for cell types. Developmental periods were stratified using BrainSpan’s general developmental ages (www.brainspan.org). Normalized expression values were scaled and plotted across cell types and developmental ages using the dotplot function of Scanpy package in Python.^62^ Expression specificity metrics were computed for individual genes using the expression weighted cell type enrichment (EWCE) method.^63^ Cell type-specific enrichment of respective network genes was estimated by using the bootstrap_enrichment_test function from EWCE R package^63^upon 10000 bootstrapping replicates. All human genes (HGNC symbols) present in the respective datasets were used as a background gene set. Transcript length and GC-content was controlled for during the analysis to avoid biases. The results were tested for multiple correction using Benjamini Hochberg method (p-adj<0.05).

### Generation of KAT-enzyme seeded co-expression networks

Post-mortem transcriptomic data of 11688 neurotypical controls was acquired from GTEx v7 database (https://www.gtexportal.org/). Sample attributes, subject phenotype and raw transcript-count files were downloaded and filtered for samples and features that were present in all brain samples and passed GTEx QC pipeline. Raw transcript counts were then normalized and log2-transformed using edgeR.^64^ Connectivity Z-scores were calculated from normalized data and all outlier samples (z-score threshold < −3) were filtered out. Further, various covariates (RINscores, age, sex, region, Hardy scale, top 13 principal components of sequencing quality-control metrics) were regressed using a linear mixed model with covariates as fixed effects and random subject intercept term. We utilized the regressed expression data from frontal cortical samples (BA9 & ACC; n=36) and constructed seeded co-expression networks for *CCBL1* (KATI), *AADAT* (KATII), *CCBL2* (KATIII), *GOT2* (KATIV), and *KMO* enzymes as described in Parikshak *et. al*.^26^ Briefly, KAT-enzyme of interest was used as a seed gene and an expanded network was generated by calculating pairwise Pearson’s correlation coefficient (PCC) between the seed gene and reference background genes (Gencode v19 annotations). For each network, genes identified as positively correlated with the seed gene (PCC>0; FDR-adjusted p<0.05) were defined as the respective KAT-positive network while genes identified as negatively correlated with the seed gene (PCC< 0; p-adj<0.05) were defined as the respective KAT-negative network. The seeded networks with top correlations (PCC > |0.5|) were visualized using igraph and ggplot2 R packages.

Co-expression gene modules for the entire dataset were computed using the robust wgcna R package,^65^in the regressed GTEx expression data.^65^ The parameters used were DS=4, MMS=1--, DCOR=0.1, PAM=False, networktype=signed, corFnc=bicor). Overall, 19 gene modules were obtained. All modules were tested for significant enrichment of top positive & negative network genes associated to each of the KAT enzymes (n=500; p-adj<0.05) using over-representation analysis (One-sided Fishers exact test). P-values were corrected using Bonferroni threshold method.

Enrichment of genes from significant modules as well as network-associated genes were assessed by performing an unbiased gene ontology (GO) analysis for GO:BP terms using gProfiler2 R package. P-values were tested for multiple correction using Benjamini Hochberg method (p-adj<0.05).

### Enrichment of schizophrenia-risk associated variants

Neurodevelopmental disease risk-associated gene sets were obtained from multiple human genetic studies. These included Hi-C defined SCZ risk genes (n=455; Mah *et al*.^37^), high confidence SCZ risk genes from GWAS (n=321; Wang *et al*.^36^), rare de novo variants associated to developmental disorders (n=285; Kaplanis et al. 2020), autism spectrum disorder (ASD)-risk genes (n= 102; Satterstrom *et al*.^38^). Enrichment of disease-risk associated gene sets in seeded networks of KAT and KMO enzymes was performed using one-sided Fishers exact test. P-values were corrected for multiple testing using Bonferroni adjustment method (*n=4*).

### Statistical Analysis

Kruskal-Wallis test with Dunn’s correction for comparisons was used to compare the calcium imaging data between treated and untreated neurons. Mann-Whitney *U* test was used to compare spine density between treated and untreated 2D neurons and organoids. For cell culture experiments, the statistical significance was analysed using GraphPad Software (version 9.5.1), California San Diego, USA (data are expressed as mean ± SEM and the statistical significance was considered when *P* < 0.05). The analyses and visualization of transcriptomic data were performed on MacOS (Ventura) in R (version 4.1) and Python (version 3.10).

### Data and Code Availability

Single-cell transcriptomic sequencing data of forebrain organoids along with the processed metadata is available at Synapse (project ID: syn26020386; https://www.synapse.org/#!Synapse:syn26020386/wiki/). All other reference single-cell sequencing data was obtained from publicly available databases. Code and intermediate data used for reproducing the transcriptomic analyses are available at www.github.com/SellgrenLab/KAT-networks. Additional information is available from the authors upon request.

## Acknowledments

We thank Eleonora Moroncina, Tintin Jansson, Tian Sang, Lihan Xu and Jessica Gracias, for technical assistance, and members of the Sellgren lab in general for helpful discussions. This study was supported by grants from Erling-Persson Family Foundation (C.M.S), the Swedish Brain Foundation (C.M.S. and M.S.;2022-0357, FO: PS2018-0058), Swedish Society for Medical Research (FO: P18-0120) and the Swedish Federal Government under the LUA/ALF agreement (C.M.S).

## Author contributions

Conceptualization, F.O., and C.M.S.; methodology, F.O., S.M.S., S.M. and C.M.S.; investigation, F.O., N.K., A.G., O.J., M.K., Š.L., L.S.; supervision, K.J., J.T., J.K. S.E., G.E., and C.M.S; writing, F.O., and C.M.S., editing, all authors.

