## Supplementary Figures for "Kynurenic acid promotes activity-dependent synaptic pruning in schizophrenia"

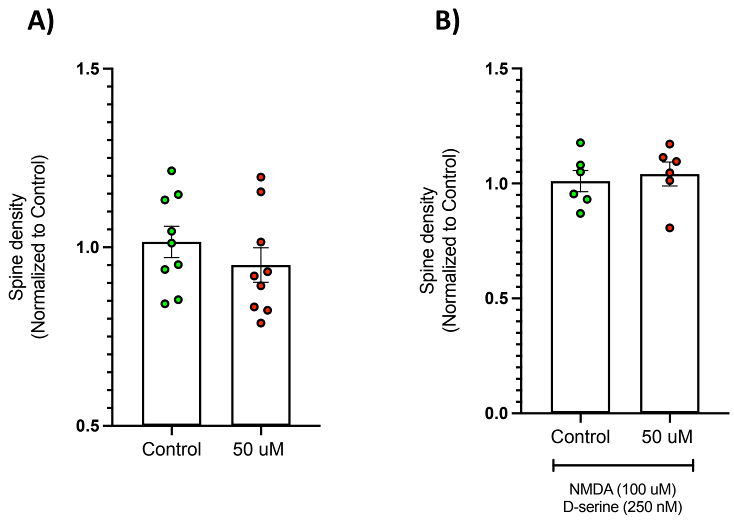


**Figure S1. Quantification of spine density in neuronal mono-cultures.** (A) Cultures treated with vehicle or 50 µM KYNA (24h). (B) Cultures treated with 50 µM KYNA or 50 µM KYNA together with 100 µM NMDA and 250 nM D-serine. Experiments were performed on day 25 neurons in duplicates and repeated three times. Data are normalized to the control group. All error bars in the figure indicate standard errors of the mean (SEM). Data was analyzed using Mann-Whitney *U* tests. All reported p-values are two-sided, statistical significance set to $p<0.05$.

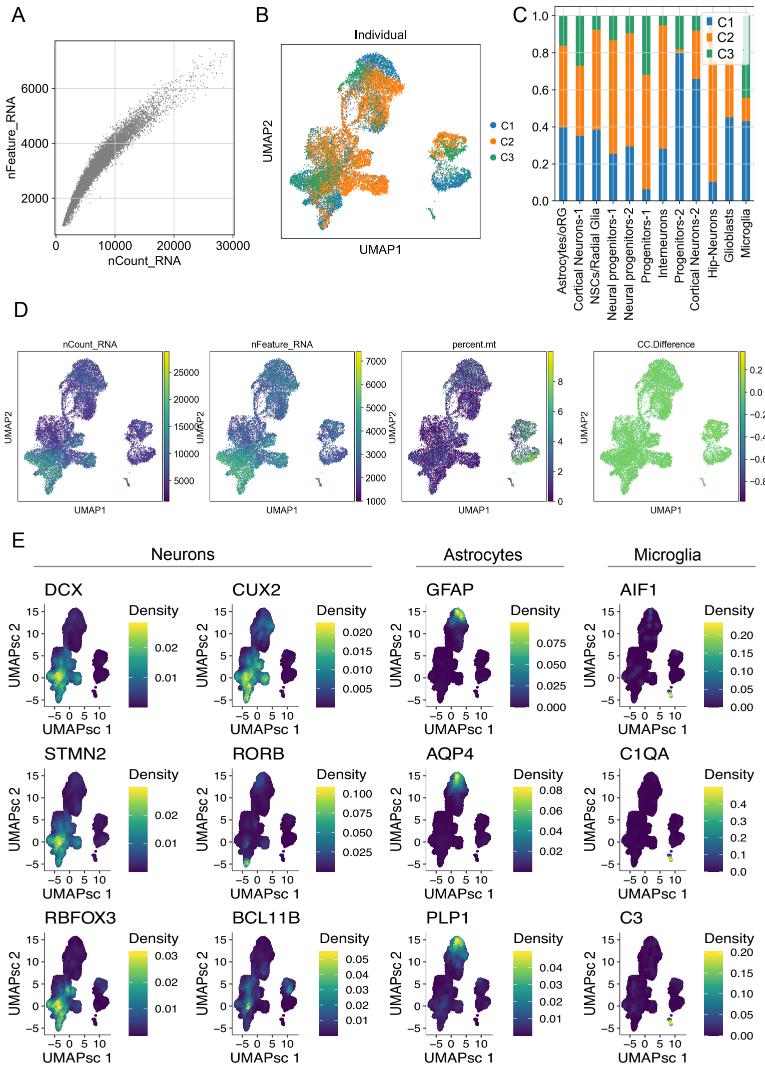

**Figure S2. Quality control of single-cell RNA sequencing of forebrain organoids.** (A) Scatter plot of all cells in the dataset with number of transcripts captured on the x-axis and number of expressed genes on the y-axis. (B) UMAP embedding of the integrated space (via BKNN method) of cells belonging to three individuals (C1,C2,C3) showing no confounding effects across clusters. (C) Bar plot of percentage of cells from the three individuals (C1,C2,C3) across cell types. (D) UMAP embedding space with cells coloured by quality control metrics post-filtering, such as number of transcripts (nCount), number of genes (nFeature), percentage of mitochondrial transcripts (percent.mt), and cell cycle score (CC.Difference). (E) Distribution of cell type-specific marker gene expression across all cells visualized on a UMAP plot. Cells are colored by their expression kernel density as shown in respective legends.

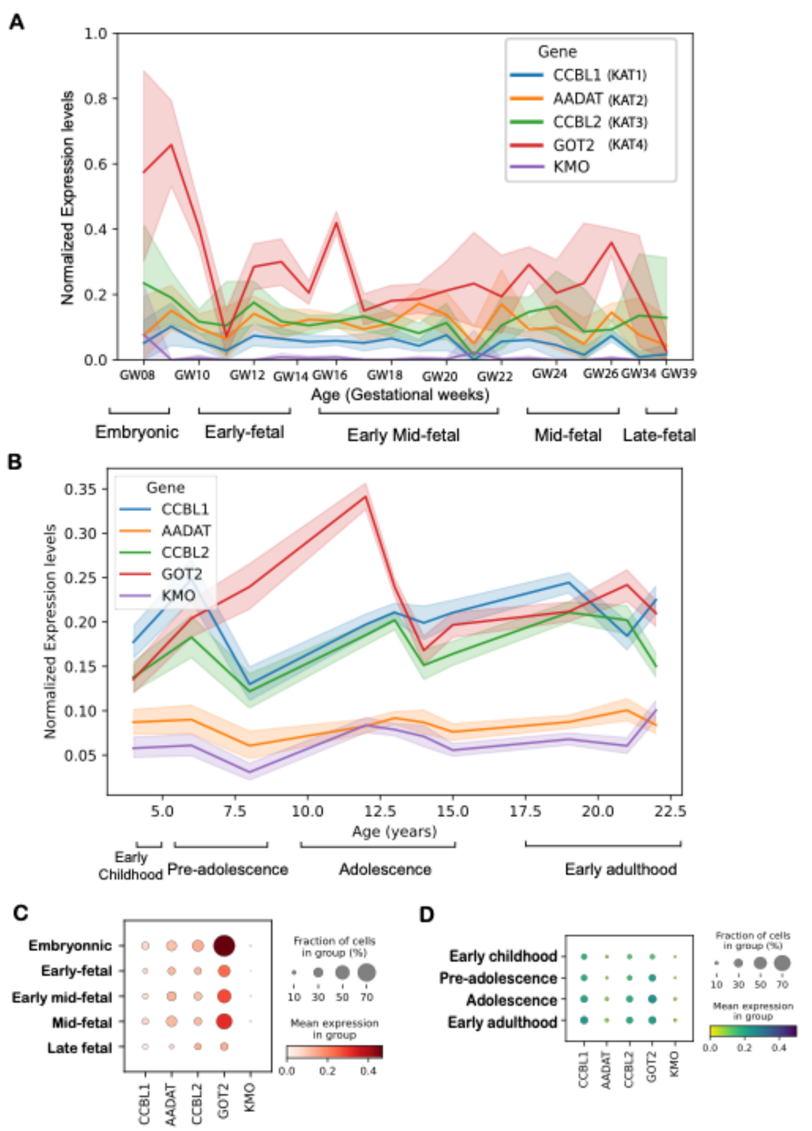


**Figure S3. Distribution of KAT-related gene expression across brain development timeline.** (A) Line chart showing the normalized expression levels (y-axis) of *KAT*-enzymes and *KMO* across fetal brain developmental age, corresponding to gestational weeks (x-axis), using single-cell transcriptomics data of human prefrontal cortex. Colored lines represent linear trends of individual genes according to the legend. 95% confidence intervals are denoted by supporting transparent lines respectively. Bottom bars display the age groups defined across the fetal developmental timeline. (B) Line chart showing the normalized expression levels (y-axis) of *KAT*-enzymes and *KMO* across postnatal brain developmental age, corresponding to years after birth (x-axis), using single-cell transcriptomics data of human prefrontal cortex.

Colored lines represent linear trends of individual genes according to the legend. 95% confidence intervals are denoted by supporting transparent lines respectively. Bottom bars display the age groups defined across the postnatal developmental timeline. Dot plot with an overview of the percentage of single cells in each age group expressing genes encoding *KAT*-enzymes, including *KMO*, as indicated by the size of the dot, during (C) fetal brain development, and (D) postnatal brain development. Color scale denotes the average normalized expression levels.

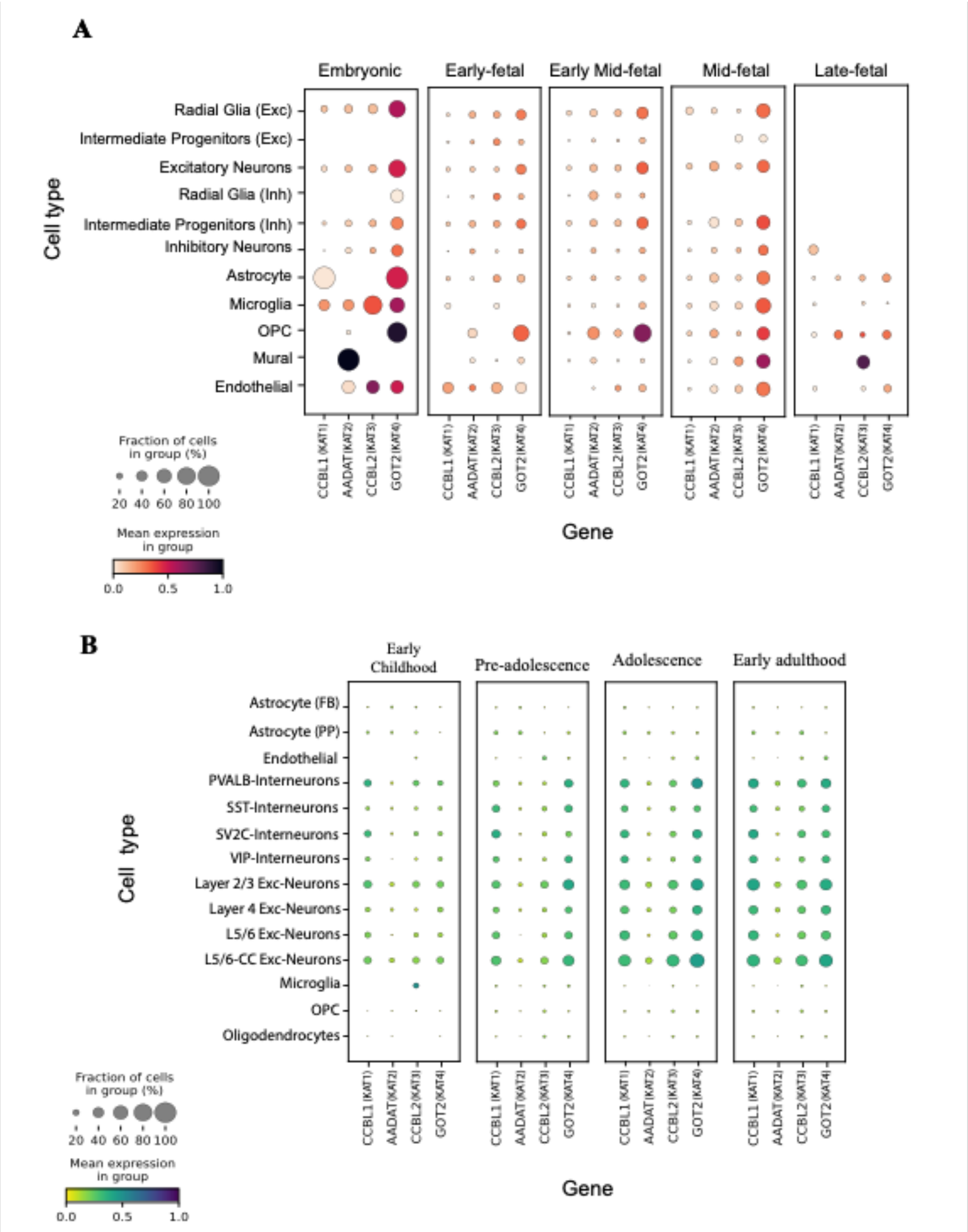


**Figure S4. Distribution of KAT-enzyme expression across cell types.** (A) Dot plot displaying the distribution of KAT-enzyme expression across cell types (y-axis) present in defined age groups during fetal brain development. [29] Size of the dots indicates the percentage of cells expressing the respective genes (x-axis) and the color scale denotes the average normalized expression. (B) Dot plot displaying the distribution of KAT-enzyme expression across cell types (y-axis) present in define age groups during postnatal brain development. Size of the dots indicates the percentage of cells expressing the respective genes (x-axis) and the color scale denotes the average normalized expression.[30]

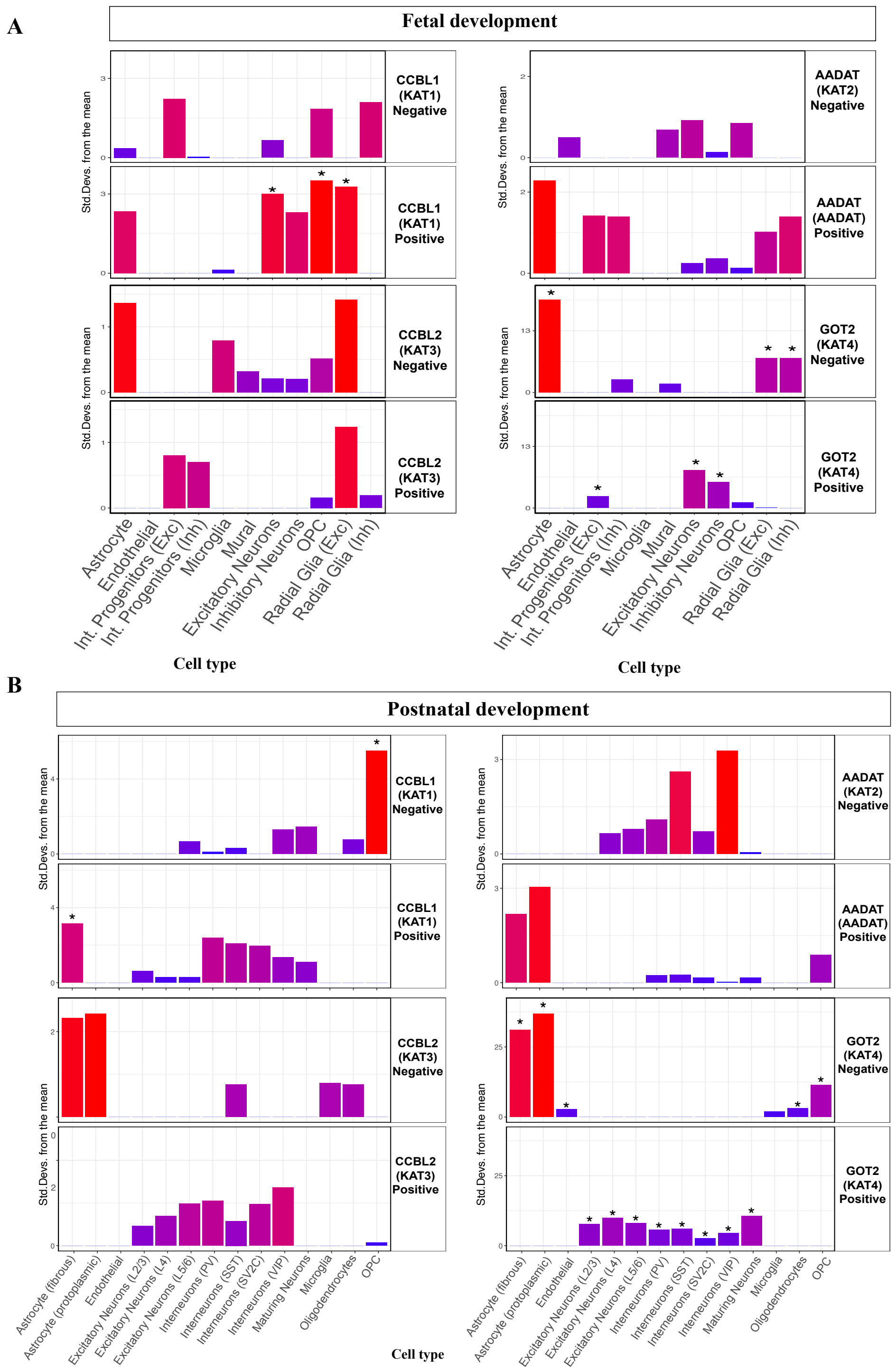


**Figure S5. Cell type-specific enrichment of genes within the *KAT*-enzyme co-expression networks.** (A) fetal brain (all age groups), and (B) postnatal brain (all age groups). Y-axis denotes the standard deviation from the mean specificity of gene set in the positive or negative networks relative to the background set (represented by the color scale). X-axis indicates the cell type group within the respective datasets. P-values were computed using bootstrapping tests (*n=10000* tests) from the Expression-weighted cell type enrichment (EWCE) method and adjusted for multiple testing using the Benjamini-Hochberg method. Asterisks denote significance at adjusted p-value<0.05. The corresponding network gene set tested for each individual plot is denoted in the right box of each bar plot.
